# In silico evidence of unique behaviors of methionine in an in-register parallel beta-sheet amyloid, suggestive of its possible contribution to strain diversity of amyloids

**DOI:** 10.1101/596742

**Authors:** Hiroki Otaki, Yuzuru Taguchi, Noriyuki Nishida

## Abstract

Mechanism of strain diversity of prions is a long-standing conundrum, because prions consist solely of abnormal isoform of prion protein (PrP^Sc^) devoid of genetic material. Pathogenic properties of prions are determined by conformations of the constituent PrP^Sc^ according to the protein-only hypothesis, and alterations to even a single residue can drastically change the properties when the residue is located at a critical position for the structure of PrP^Sc^. Interestingly, methionine (Met) is often recognized as the polymorphic or species-specific residues responsible for species/strain barriers of prions, implying its unique influences on the structures of PrP^Sc^. However, how it is unique is difficult to demonstrate due to lack of the detailed structures of PrP^Sc^. Here we analyzed influences of Met substitutions on structures of an in-register parallel β-sheet amyloid of α-synuclein (αSyn) by molecular dynamics (MD) simulation, to extrapolate the results to PrP^Sc^. The MD simulation revealed that Met uniquely stabilized a U-shaped β-arch of the Greek-key αSyn amyloid, whereas other hydrophobic amino acids tended to destabilize it. The stabilizing effect of Met was attributable to the long side chain without Cβ branching. Our findings exemplify specifically how and in what structure of an in-register parallel β-sheet amyloid Met can uniquely behave, and are suggestive of its influences on structures of PrP^Sc^ and strain diversity of prions. We also discuss about relations between α-helix propensity and local structures of in-register parallel amyloids.

## Introduction

Prion diseases are a group of neurodegenerative disorders which are characterized by accumulation of the abnormal isoform (PrP^Sc^) of prion protein (PrP) in the central nervous system [1]. Prion diseases have three etiologies, i.e., sporadic, inherited and acquired, depending on how the causative PrP^Sc^ starts propagation in the body; in sporadic and inherited prion diseases, e.g., sporadic Creutzfeldt-Jakob disease (CJD) and fatal familial insomnia (FFI), respectively, the causative PrP^Sc^ are generated by spontaneous conformational conversion of endogenous normal isoform PrP into PrP^Sc^ without or with mutation in PRNP gene, which encodes PrP. Acquired prion diseases are caused by intake of exogenous PrP^Sc^ as infectious agents, prions, e.g. epidemic bovine spongiform encephalopathy (BSE) in cattle [2] and chronic wasting disease (CWD) in cervids [3]. Prions behave similarly as viruses, with high infectivity, existence of many strains, species/strain barriers, and adaptation to new hosts, despite the lack of conventional genetic material. These virus-like properties of prions are explained by hypothesis that pathogenic properties of prions are enciphered in the conformations of PrP^Sc^, i.e., protein-only hypothesis [1][4].

Indeed, pathogenic properties of prions and the consequent clinical features are greatly affected by the primary structure of constituent PrP^Sc^ even by alteration of a single residue, in accordance with the fundamental fact that conformations of proteins are determined by the primary structures. For example, in sporadic CJD, PrP deposition patterns, lesion profiles in the brain, clinical presentations, and apparent molecular sizes of protease-resistant cores of PrP^Sc^ are varied depending on whether the polymorphic codon 129 is methionine (Met) or valine (Val) [5]. A pathogenic mutation D178N of human PrP reportedly causes either FFI or familial CJD in association with Met129 or Val129, respectively [6]. The codon 129 polymorphism also determines susceptibility to prion infections, i.e., species/strain barriers: a vast majority of new-variant CJD cases which resulted from trans-species transmission of BSE to humans are homozygous for Met129 [7][8]; the polymorphic codon 132 of elk PrP, either Met or leucine (Leu), which is equivalent to the codon 129 polymorphism of human PrP, also affects susceptibility to CWD [9][10]. From the viewpoint of the protein-only hypothesis, effects of the polymorphisms are mediated by structural alterations of the constituent PrP^Sc^. However, specifically how the subtle differences between the hydrophobic amino acids affect structures of the PrP^Sc^, to an extent to drastically affect susceptibility and clinicopathological features, is not identified yet. Detailed structures of PrP^Sc^ are necessary for the investigation but they are unavailable due to its incompatibility with conventional high-resolution structural analyses. Based on available data, PrP^Sc^ is currently hypothesized either an in-register parallel β-sheet amyloid or a β-solenoid [11][12][13][14][15][16].

Remarkably, not only at the polymorphic codon 129, Met at different positions of PrP are often involved in strain diversity and species barriers of prions. In amyloid formation reactions among Y145Stop of human PrP and the equivalent mutants of mouse and Syrian hamster PrPs, Met at residues 137-138 (in human numbering throughout, unless otherwise noted) are critical for efficient amyloid formation and cross-seeding among them [17][18]. Ile137Met substitution substantially extended incubation periods of experimental transmission of prions from sporadic CJD homozygous for Met129 [19]. Met109 and Met111 are influential on transmission efficiencies among different hamster species [20][21]. In an experiment observing influences of systematically-introduced mutations of mouse PrP on conversion efficiencies in scrapie-infected Neuro2a cells, Met at some positions, e.g., Met213, were replaceable with other hydrophobic residues like Leu or isoleucine (Ile), whereas unreplaceable at other residues, e.g., Met206 [22]. In cross-seeding between ovine and cervine PrPs, I208M (in ovine numbering) mutation showed profound influences on seeding efficiencies [23]. Given those documented facts, we asked ourselves whether Met could have distinct properties than other hydrophobic amino acids that uniquely affect structures of PrP^Sc^. To address the question, we exploited an in-register parallel β-sheet amyloid of α-synuclein (αSyn) as a surrogate structural model for PrP^Sc^, as we previously did [24][25].

αSyn amyloid is the main component of Lewy body, which is a hallmark of Parkinson’s disease (PD) and dementia with Lewy bodies (DLB), and is known to have prion-like properties including transmissibility and strain diversity [26]. For instance, αSyn forms various types of amyloids in vitro which are different in appearances, proteolytic fragment patterns, and cytotoxicity [27]. Moreover, αSyn amyloids isolated from DLB and multiple system atrophy have different proteolytic fragment patterns, suggestive of distinct conformations [28]. Those are highly reminiscent of prions and imply that prion-like properties are inherent in in-register parallel β-sheet structures. Unlike PrP^Sc^, detailed structures of αSyn amyloids have been determined by solid-state NMR (ssNMR) [29] or cryo-electron microscopy [30][31], and we used a Greek-key αSyn amyloid (PDB ID: 2n0a [29]) to assess whether Met behaves uniquely in the in-register parallel β-sheet structures, particularly in terms of effects on stability of amyloid structures. As a result, our study revealed unique effects of Met in a U-shaped β-arch of the amyloid that could be attributed to the long side chain without Cβ branching. If PrP^Sc^ is also an in-register parallel β-sheet amyloid, the influences of Met on structural stabilities of amyloid may contribute to a mechanism underlying the strain diversity of prions.

## Results

### Predicted intrinsic propensities of **α**Syn(E61M) and **α**Syn(G84M)

To compare influences of Met and other hydrophobic amino acids on the local structures of αSyn amyloids, we introduced Met substitution at residues 61 (E61M) and 84 (G84M), where we previously investigated influences of isoleucine substitutions [24]. We first estimated intrinsic propensities of the mutant αSyn by secondary structure prediction algorithm [24][32][33]. They both raised local (Pα-Pc) values and the positive-(Pα-Pc) areas coalesced across the loop regions (Fig 1A, **red circles, and** 1B). Notably, in the mutant αSyn with G84M, αSyn(G84M), the positive-value areas of (Pβ-Pα) and (Pβ-Pc) in residues 88-90 disappeared, making the C-terminal region solely (Pα-Pc)-positive. Those predicted propensity profiles of αSyn(E61M) and αSyn(G84M) were partly anticipated because Met has high α-helix propensity due to its long and non-branching side chain [34][35]. Given the raised helix propensities, we initially thought that they might destabilize the loops.

**Fig 1.**
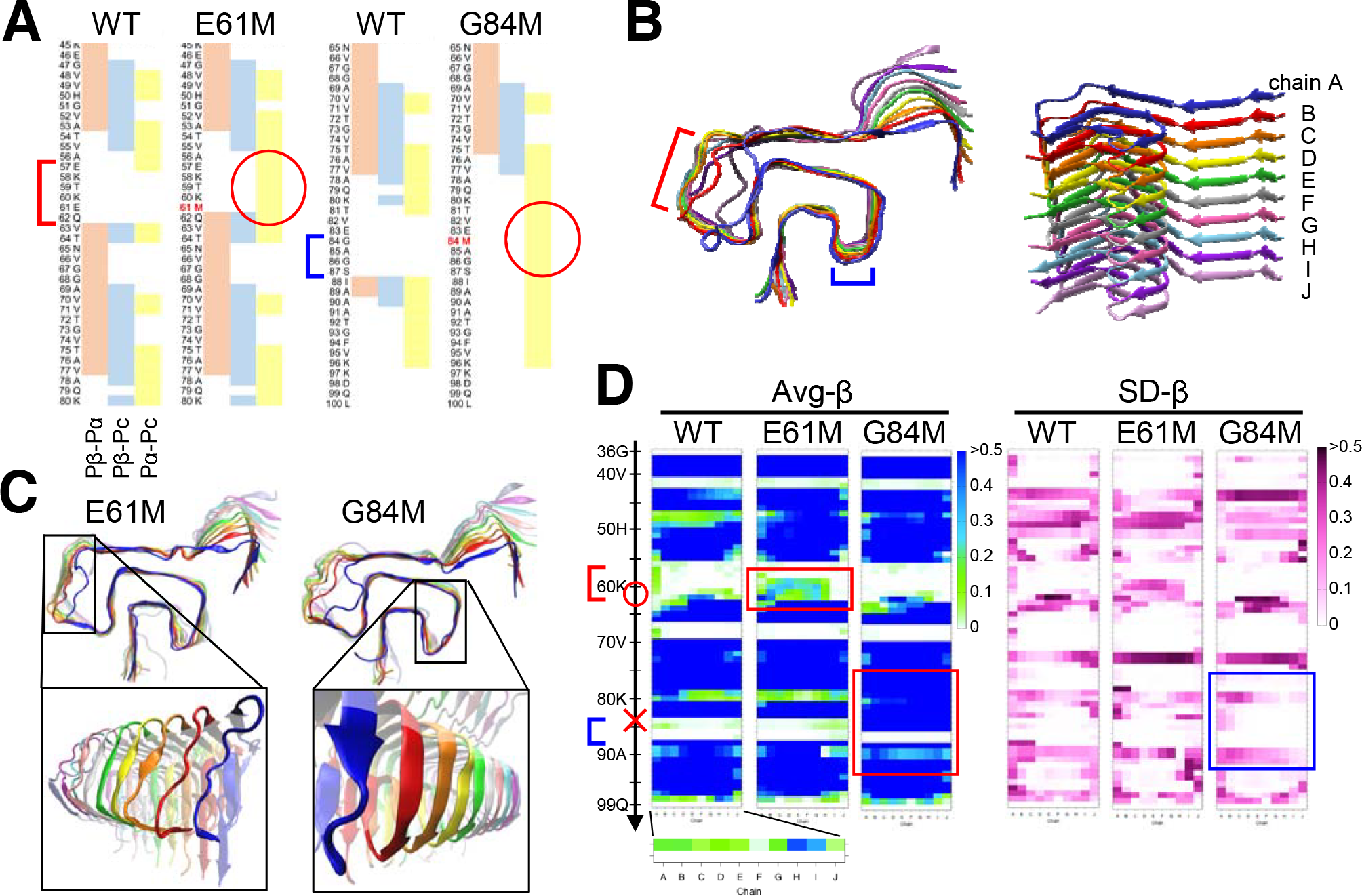
Influences of methionine (Met) substitutions on local structures of the “Greek-key” αSyn amyloid. **A.** Diagrams of secondary structure predictions profiles of αSyn(E61M) and αSyn(G84M), showing the magnitude relations between Pα, Pβ, and Pc, i.e., Pβ-Pα, Pβ-Pc, and Pα-Pc; residues with zero (“0”) or positive values are indicated in red, blue and yellow, respectively. E61M and G84M substitutions are indicated in red letters. WT, wild-type αSyn. Red and blue brackets indicate positions of loop(57-62) and loop(84-87), respectively. **B.** A final snapshot of αSyn(WT) after 400 ns of MD simulation of a “Greek-key” αSyn amyloid (PDB ID: 2n0a) (left). Red and blue brackets correspond to the loops in Fig 1A. Chains of the amyloid stack are referred to as chain A-J from front to back (right). **C.** Final snapshots of αSyn(E61M) (left) and αSyn(G84M) (right) amyloids after 400 ns of MD simulations. The insets view the indicated regions from slightly different angles to show structures of each chain. Note that β-sheets are occasionally induced in loop(57-62) of αSyn(E61M) amyloid (left inset) and, in αSyn(G84M) amyloid, loop(84-87) is converted to a stable bent β-sheet (right inset). **D.** Heatmaps of the average β-sheet propensity values (Avg-β) and the standard deviations of the values (SD-β) based on five (for WT) and three (for the mutant) independent runs of 400-ns MD simulations. Vertical axes and horizontal axes of all the heatmaps represent the residue numbers and the chains A-J, respectively. Red circle and cross on the vertical axis indicates positions of E61M and G84M, respectively. Avg-β values in loop(57-62) of αSyn(E61M) and in loop(84-87) of αSyn(G84M) amyloids were increased (red boxes) corresponding to the induced β-sheets in those regions (Fig 1B). The relatively low SD-β values in αSyn(G84M) amyloid (blue box) reflect the stabilized local structures.

### MD simulation of αSyn(E61M) and αSyn(G84M)

MD simulations of homo-oligomer amyloids of αSyn(E61M) and αSyn(G84M) unexpectedly revealed that both the mutations tended to stabilize the local structures particularly when they induced new β-sheets in the nearby loops (Fig 1C). E61M substitution occasionally induced β-sheets in the loop encompassing rsidues 57to 62, loop(57-62) (Fig 1C, **left inset**) and, once it occurred, the local structure was stabilized. The induction of β-sheets were mainly in chains C-H and the loops of chains A-B tended to disorder (Fig 1C, **left inset, and** Fig 1D, **red box**), unlike αSyn(E61I) amyloid whose induced β-sheet were mainly on the chain-A side [24]. Effects of G84M substitution were more impressive. The almost entire region C-terminal to residue 80, even including the loop encompassing residues 84 to 87, loop(84-87), reproducibly converted to stable β-sheets (Fig 1C, **right inset**) and the entire molecule was stabilized (Fig 1D, **red and blue boxes of G84M**), in contrast to those of G84I substitution [24]. Incidentally, we assessed behaviors of αSyn(G84L) amyloid in MD simulation, which showed a similar predicted propensity profiles as αSyn(G84M) with positive (Pα-Pc) values across loop(84-87) (Fig 2A), consistent with comparable α-helix propensity of Leu to Met. Unexpectedly, the loop region was substantially destabilized in αSyn(G84L) amyloid like that of αSyn(G84I) [24] (Fig 2B, **red circle, and** 2C, **red and blue boxes**). Being interested in the stabilizing effects of G84M and the differences from G84L, we focused on the residue 84 and performed further in silico analyses to identify the underlying mechanisms.

**Fig 2.**
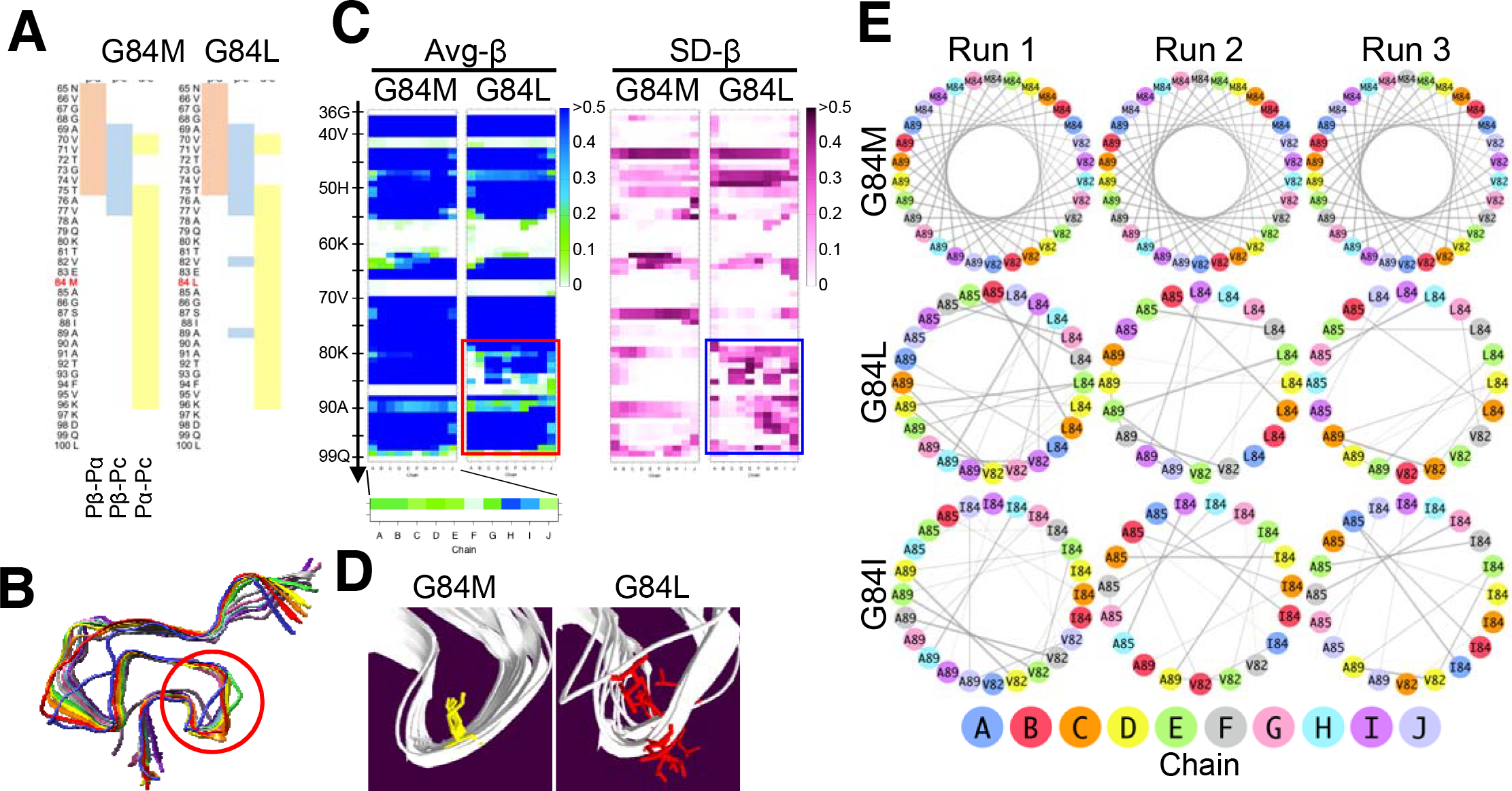
Influences of G84L and G84M are rather different, despite the absence of Cβ-branching in common. **A.** Diagram of secondary structure prediction profiles of αSyn(G84L) in comparison with αSyn(G84M). **B.** A final snapshot of αSyn(G84L) amyloid after 400 ns of MD simulation. The local structures around the mutation were disturbed (red circle), unlike the αSyn(G84M) amyloid. **C.** Heat maps of Avg-β and SD-β of αSyn(G84L) amyloid based on three independent runs of 400 ns. The structural disturbances are reflected in both (red and blue boxes, respectively), in contrast with αSyn(G84M) amyloids. **D.** Directions of the side chains of residue 84 after 400 ns of simulation. Some side chains of Leu84 are pointing outward, while those of Met84 stay inside. **E.** Hydrophobic contact diagrams based on simulation of 400 ns. Only the residues involved in the interactions with residue 84 are presented. Residue numbers and chains are indicated by numbers and colors of the dots, respectively. Thickness of each line corresponds to occupancy of the hydrophobic contact.

### Further analyses of **α**Syn with substitution at the residue 84

We noticed that the final status of 400 ns simulations were different between αSyn(G84M) and αSyn(G84L) amyloids; all the side chains of Met84 pointed inward of loop(84-87), whereas some side chains of Leu84 flipped and pointed outward (Fig 2D). Hydrophobic contact diagrams of homo-oligomer αSyn(G84M) and αSyn(G84L) amyloids clearly illustrated the differences: Met84 of αSyn(G84M) amyloid had intra-chain interactions evenly with residues 82 and 89 of the same chain in all the layers, and the interactions were rather reproducible (Fig 2E, **G84M, Run 1-3**); on the other hand, αSyn(G84L) amyloids only infrequently showed that pattern of interactions (Fig 2E, **G84L**) with frequent intra- or inter-chain interactions between Leu84 and either residue 82 or 89 that were hardly reproducible (Fig 2E, **G84L, Run 1-3**). The results were similar to those of another destabilizing mutation G84I (Fig 2E, **G84I**), which we reported elsewhere [24].

We noticed that the flipping of side chains of Ile84 occurs as early as immediately after energy minimization (Supplementary Fig S1A). Even before energy minimization, side chains of Leu84 and Ile84 were placed toward the periphery of the β-arches by the software SCWRL4, apparently inclined to flip (Fig 3A, **Before**), possibly due to narrow inner spaces of the β-arches for the bulky side chain of Leu or Ile. In contrast, Met84 wedged deeply into the β-arches (Fig 3A, **G84M, Before**) and maintained the positions also after energy minimization (Fig 3A, **After**).

**Fig 3.**
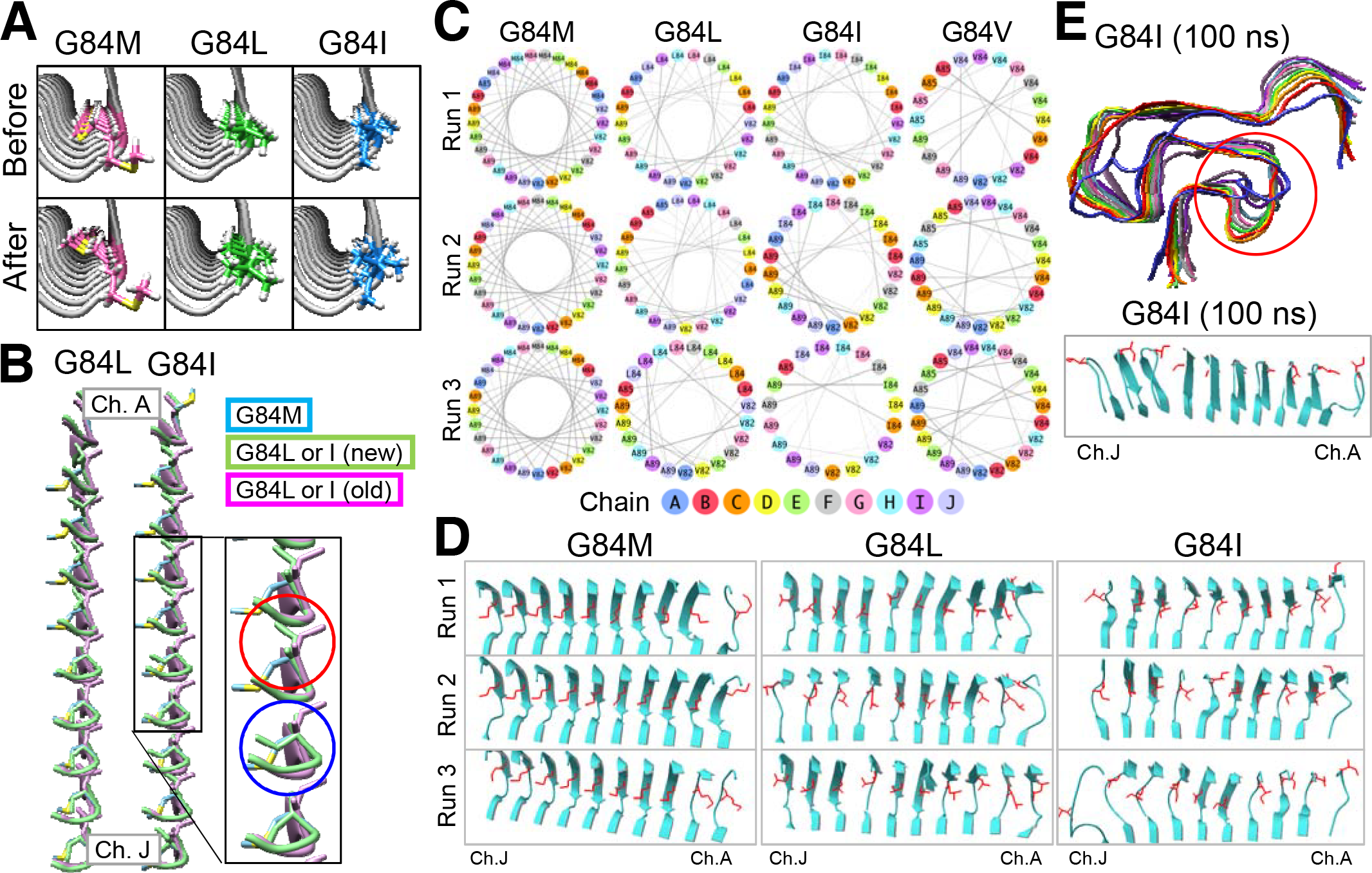
The varied structural disturbances in MD simulation among αSyn(G84M), αSyn(G84L) and αSyn(G84I) amyloids are not due to differences in their initial status. **A.**Comparison of status of the three mutant αSyn amyloids before and after energy-minimization step. **B.**Side views of the status of side chains of Leu84 and Ile84 after modeling of the mutant αSyn amyloids by a software SCWRL4, comparing a modeling based on the αSyn(WT) (“G84L or I (old)”, purple chains) and another based on the energy-minimized αSyn(G84M) amyloid (“G84L or I (new)”, green chains). Only residues 83-85 are shown. **C.**Hydrophobic contact diagrams of mutant αSyn amyloids based on short MD simulations of 5 ns, after modeling based on the minimized αSyn(G84M) amyloid. Only the residues involved in the interactions with residue 84 are presented. **D.**Final snapshots of the mutant αSyn amyloids of the same short-MD simulations as in Fig 3C, presenting directions of the side chains of residues 84 in red. Only the region from Lys80 to Ala90 is shown. **E.** A final snapshot of the MD simulation of αSyn(G84I) amyloid where simulation of “Run 1” in Fig 3D was extended up to 100 ns (upper panel), and status of the side chains of Ile84 (lower panel). The loop region was viewed from outside of the β-arch to show flipping of the side chains.

To make the initial status same, we created another set of αSyn(G84L) and αSyn(G84I) amyloid models by replacing Met84 residues of an energy-minimized αSyn(G84M) amyloid (Fig 3A, **“After” of G84M**) with Leu or Ile, and subjected them to MD simulations (Supplementary Fig S1B). After the remodeling by SCWRL4 based on the energy-minimized αSyn(G84M), the side chains of Leu84 or Ile84 of chains A-E still similarly positioned as in the old modeling (Fig 3B, **red circle**) and those of chains F-J maintained the same initial position as those of αSyn(G84M) (Fig 3B, **blue circle**). In the following MD simulation, only in 5 ns, differences between αSyn(G84M), αSyn(G84I) and αSyn(G84L) amyloids were more accentuated, and they behaved similarly as in the 400-ns simulations, as reflected in hydrophobic contact diagrams (Fig 3C), i.e., Met84 constantly interacting with residues 82 and 89 while others showing unstable interactions. Final status of side chains after the 5-ns runs substantially differed between αSyn(G84M) and αSyn(G84L) or αSyn(G84I) (Fig 3D). While the side chains of Met84 maintained the same positions and directions as in the initial status except for those of chains A and B (Fig 3D, **G84M**), those of Leu84 and Ile84 pointed different directions with some flipping and local β-sheet structures were lost (Fig 3D, **G84L and G84I**). Although disturbances of the loop structures were relatively small in Run 1 of the 5-ns MD simulation of αSyn(G84I) (Fig 3D, **G84I, Run 1**), in the following 100 ns, the loop and adjacent structures substantially disordered (Fig 3E, **red circle**) and some side chains flipped (Fig 3E, **lower panel**). We also compared αSyn(G84V) amyloid because Val has relatively short side chain with Cβ branching, in contrast with the side chain of Met. As expected, Val84 behaved very differently from Met84 as reflected in the hydrophobic contact diagrams (Fig 3C, **G84V**).

Collectively, the long and unbranched side chain of Met84 could be responsible for the unique stabilizing effects on the β-arch. The absence of Cβ branching in the side chain allows Met to behave distinctly from Ile and Val. Leu has an unbranched side chain but possibly the length is not sufficient to reach Ala89 for hydrophobic-core formation to stabilize the β-arch.

### Influences of G84F and G84A, which both lack C**β** branching but are different in lengths of side chains

To assess influences of length of side chains on the local structures, we performed short MD simulations of amyloids of αSyn(G84F) with phenylalanine (Phe) and αSyn(G84A) with alanine (Ala) at residue 84. Length of the side chain of Phe is slightly shorter than that of Met but longer than that of Leu; Ala has the shortest side chain except for glycine. Because of the moderate to high α-helix propensities of Ala and Phe, their secondary structure prediction profiles were similar to that of αSyn(G84M) albeit a gap in the positive-(Pα-Pc) area across the loop (Fig 4A). In the MD simulations, disturbance of the backbone structures by G84A substitution was limited, with small derangement in the patterns of heat map of Avg-β over 5 ns (Fig 4B, **G84A**). Although side chains of Ala84 tended to flip (Fig 4C), steric effects of the small side chain were possibly tolerable. In the αSyn(G84F) amyloid, a certain degree of destabilization of the local structures was suggested by the gradual decrements in Avg-β values over 5 ns, particularly on the chain-J side (Fig 4B, **G84F**). Regardless, the side chains of Phe84 were relatively maintained in the original positions without flipping and most of the phenyl groups are constantly located near Ala89, although their bulkiness seemed to disturb the surrounding structures (Fig 4D **and Supplementary Movie, G84F**).

**Fig 4.**
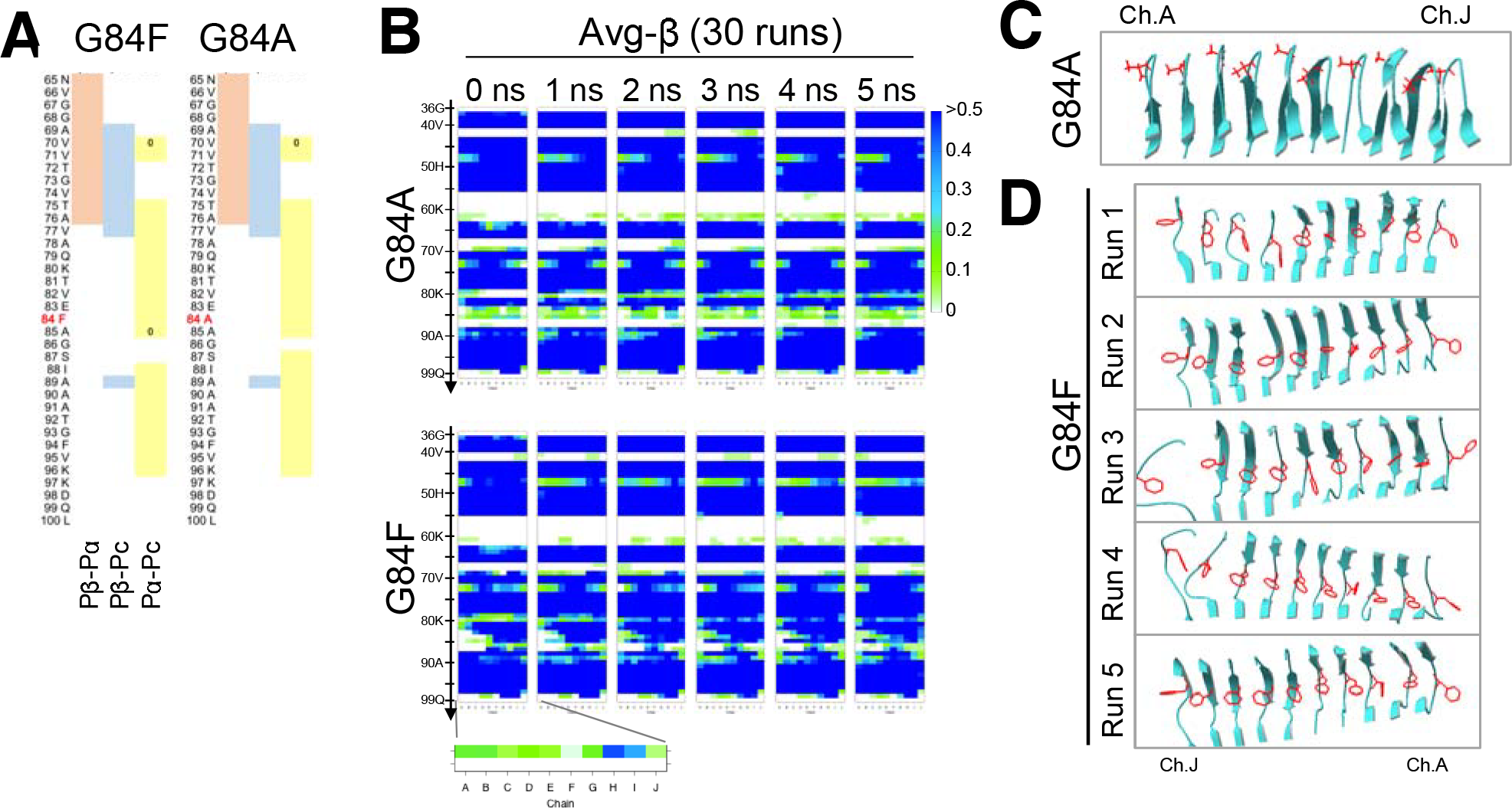
The absence of Cβ branching and lengths of the side chains are important for compatibility with the U-shaped β-arch. **A.** Diagram of secondary structure prediction profiles of αSyn(G84F) and αSyn(G84A). **B.** Heat maps of Avg-β of αSyn(G84A) and αSyn(G84F) amyloids based on 30 independent MD runs of 5 ns. **C.** Directions of the side chains of Ala84 after 5 ns of simulation in Run 1 of the 30 independent 5-ns MD simulation runs. The loop region was viewed from outside of the β-arch, to show flipping of side chains. Only the region from Lys80 to Ala90 is shown. **D.** Directions of the side chains of Phe84 after 5 ns of simulation in the first five runs of the 30 independent 5-ns MD simulations.

## Discussion

The series of MD simulation of in-register parallel β-sheet amyloids of αSyn demonstrated distinctive influences of Met on local structures of the amyloid among other hydrophobic amino acids. The uniqueness was conformation-dependent: Met84 outstandingly stabilized the U-shaped loop(84-87) while Ile, Leu and Val at residue 84 destabilized the structures. On the other hand, Met61 showed similar influences on the large loop(57-62) as Ile61 albeit subtle differences in terms of the layers where β-sheets were induced. The stabilization of the loop region by Met61 and Ile61 is attributable to elimination of the negative charge of the original glutamate at residue 61 that can destabilize the local structures by repulsion between one another and also to the hydrophobic side chains of Met and Ile that contribute to hydrophobic cores of the β-arches. Cβ-branching of the side chain of Ile would not be a problem in the position.

In contrast, the Cβ-branching side chain of Ile84 or Val84 can destabilize U-shaped loops like loop(84-87). Steric clash of the bulky Cβ-branching side chains on the bent backbones and other side chains of β-arches are disadvantageous for maintaining the conformation, as demonstrated in αSyn(G84I) and αSyn(G84V) amyloids. In such structures, Met would behave distinctly from those amino acids because of the absence of Cβ branching and, moreover, because of the longest side chain among hydrophobic amino acids except for tryptophan. In the MD simulation of αSyn(G84M) amyloid, the ends of side chains of Met84 were fixed in the vicinity of Ala89 across the inner space of the β-arch (Fig 2E, **G84M, and Supplementary Movies, G84M**), stabilizing the β-arch like a “crossbeam”. Our studies also demonstrated that Leu cannot necessarily substitute for Met despite their commonality in terms of hydrophobicity and non-branching Cβ, when stable hydrophobic interactions with Ala89 five-residues away was required. Not only in the U-shaped β-arch of αSyn amyloid, the long hydrophobic residue of Met might be likewise essential in maintaining a stable hydrophobic core of β-arch in a region with sparse hydrophobic residues a few residues apart from each other. In reality, even subtle difference between Met and Leu are significant in certain cases of prion propagation, as Met/Leu polymorphism of elk PrP affects susceptibility to CWD infection [9][10]. If the sequence comprising residue 132 (in elk numbering), V_125_GGL_128_GGY(M/L)_132_LGSA_136_MS, forms a β-arch with the side chain of residue 132 directed inward, the long side chain of Met could be more favorable for hydrophobic-core formation, because Leu128 and Val125 are four- and seven-residue away, respectively. Given the effects of Cβ-branching and length of side chains, Val and Met should cause very different influences on local structures in certain β-arches; possibly their different behavior may explain how Met/Val polymorphisms at residue 129 of human PrP affect clinicopathological features and susceptibilities to certain prion strains, e.g., new-variant CJD [7][8].

We propound this type of contribution of Met to structural stability of amyloids as just one possible model. As the long hydrophobic side chain of Met would be also advantageous for efficient inter-domain interactions and inter-molecular interactions, as seen in an Aβ amyloid (PDB ID: 2mpz) [36], possibility of other models need to be considered as well.

The secondary structure prediction algorithm may fortuitously predict β-sheets and loops of in-register parallel β-sheet amyloids to some degree [24], because it conveniently assigns high Pα values to Ala and residues with long side chains without Cβ branching, e.g., Met, Leu and glutamine, while assigning high Pβ values to residues with intermediate-length side chains with Cβ branching, e.g., Val, Ile and threonine. If residues without Cβ-branching could flexibly contribute to structural stabilization of various β-arches including U-shaped ones as suggested by the present study, a region of high α-helix propensity with Met or Leu may be actually advantageous for amyloid formation because it can adapt to various forms of β-arches. Even though such a region may resist to conversion from α-helix form to amyloid form due to requirement for the energy to unfold, it can contribute to amyloid formation by stabilizing β-arches once successfully converted. Hydrophilic residues with α-helix propensity like glutamine can also contribute to stabilization of amyloid structures by making “glutamine ladders” [37]. Prediction algorithms optimized for specific use on amyloids are desirable because the algorithm used here tend to assign inappropriately high coil propensities to a proline residue than its actual effects on amyloid formation.

A caveat to the present study is that the setting was very artificial, i.e., starting with mutant αSyn perfectly fit to the Greek-key conformation irrespective of mutant types and suddenly unleashing them to behave according to thermodynamic principles. In reality, higher freedom of motion of the peptide backbone and potential steric effects of the long side chain of Met would pose energy barrier for Met to settle on β-arches. Indeed, spontaneous amyloid formation by Y145Stop mutant of Syrian-hamster PrP which has more Met residues than mouse or human PrP counterparts was inefficient [21]. Such inefficiency in refolding can divert a certain fraction of molecules to aberrant refolding pathways in in-vitro or in-vivo experiments to complicate interpretation of the results. Regardless of those limitations, MD simulation provides a direct insight into structures and behaviors of in-register parallel β-sheet amyloids, and this approach can shed light on the mechanisms of strain diversity of amyloids including PrP^Sc^.

In conclusion, Met can uniquely behave in certain structures of in-register parallel β-sheet amyloids because of the long side chain without Cβ branching, and might be essential in stabilizing certain U-shaped β-arches like loop(84-87) of αSyn amyloid. The effects on local structures might underlie sequence-dependent species/strain barriers when the site is an interface between substrate and template amyloid where the substrate initiate the conversion.

## Materials and Methods

Detailed protocols of the modeling mutant αSyn amyloids, MD simulation, short MD simulation, and various analyses were mostly the same as described in the preprint on bioRxiv [25]. Here we briefly describe essential points.

### Secondary structure prediction

We used the same algorithm (http://cib.cf.ocha.ac.jp/bitool/MIX/) [32] as previously reported [24][33]. To clarify magnitude relations between two of the three conventional parameters, i.e., α-helix propensity (Pα), β-sheet propensity (Pβ), and coil propensity (Pc), we used their arithmetic differences as a new set of parameters, i.e., (Pβ-Pα), (Pβ-Pc) and (Pα-Pc) [24].

### Modelling Structure

We used the Greek-key αSyn amyloid (PDB ID: 2n0a [29]) as the starting structure, after truncating the disordered N- and C-terminal regions (residues 1–35 and 100–140, respectively) [25]. The N- and C-termini were acetylated and *N*-methylated using AmberTools16 [38]. For modelling αSyn mutants, we used SCWRL4 [39]. Modelled amyloids were solvated with a rhombic dodecahedron water box with a minimum wall distance of 12 Å using GROMACS (version 5.0.4). Na+ and Cl– ions were randomly placed to neutralize the system and yield a net concentration of 150 mM NaCl. The protonation state of the histidine at the residue 50 was fixed as the Nδ-H tautomer (HID form in AMBER naming convention) in all simulations.

### MD simulations

GROMACS (versions 5.0.4 and 5.1.2) [40] with the AMBER ff99SB-ILDN force field [41] was used for MD simulations with the TIP3P water model [42]. The system was initially minimized for 5,000 steps with the steepest descent method, followed by 2,000 steps with the conjugate gradient method. During minimization, heavy atoms of the amyloids were restrained with a harmonic potential with a force constant of 10.0 kcal/mol•Å2. After minimization, the temperature of the system was increased from 0 to 310 K during a 1 ns simulation with the restraints. Next, a 1 ns equilibration run was performed by gradually reducing the restraints from 10.0 kcal/mol•Å2 to zero and subsequent equilibration was performed in the NPT ensemble for 2 ns at 310 K and 1 bar. Production runs were carried out for 400 ns in the NPT ensemble at 310 K and 1 bar (Supplementary Figure S1C). We used the velocity-rescaling scheme [43] and the Berendsen scheme [44] for thermostat and barostat, respectively. The LINCS algorithm [45] was used to constrain all bonds with hydrogen atoms, allowing the use of a 2 fs time step. Electrostatic interactions were calculated with the Particle-mesh Ewald method [46]. The cut-off length was set to 12 Å for Coulomb and van der Waals interactions. The Verlet cut-off scheme [47] was used for neighbour searching. Trajectory snapshots were saved every 10 ps for the MD simulations.

### Analyses

As for analyses of 400-ns simulations, the first 100 ns of each MD trajectory was discarded, and 30,001 snapshots (trajectory from 100 to 400 ns) were used. The secondary structure content during the simulations was calculated using DSSP [48,49] and gmx do_dssp in GROMACS. Hydrophobic contacts were analysed using PyInteraph [50]. A hydrophobic contact was assigned if the distance between the centres of mass of side chains is less than 5 Å [51]. The results of hydrophobic contact analyses were visualized using Cytoscape (version 3.5.1) [52]. All molecular structures for αSyn amyloids were drawn by UCSF Chimera (version 1.12) [53] and Swiss PDB Viewer [54]. The movies of trajectories of MD simulations were prepared with VMD [55].

### Short MD runs

We carried out 5 ns production runs from the common minimized structure according to the almost same procedure as the 400 ns MD simulations. For the simulation of αSyn(G84L), αSyn(G84I) and αSyn(G84V) amyloids, the initial structures were modeled by replacing Met84 of an energy-minimized αSyn(G84M) amyloid by using SCWRL4 [39] (Supplementary Figure S1C). We carried out 5 MD simulations for αSyn(G84M), αSyn(G84L), αSyn(G84I) and αSyn(G84V), and 30 MD simulations for αSyn(G84F) and αSyn(G84A).

## Acknowledgement

We thank Kei Yura, Ochanomizu University, for generously allowing us to use the neural network secondary structure prediction algorithm on his website. The numerical calculations were carried out on the TSUBAME2.5/3.0 supercomputer at the Tokyo Institute of Technology and the Reedbush-U supercomputer at the Information Technology Center, the University of Tokyo. This work was supported by the “TSUBAME Encouragement Program for Young/Female Users” of Global Scientific Information and Computing Center at the Tokyo Institute of Technology, the “Initiative on Promotion of Supercomputing for Young or Women Researchers” from the Information Technology Center, the University of Tokyo, and Takeda Science Foundation (www.takeda-sci.or.jp/).

---

**Supplementary Figure S1.**
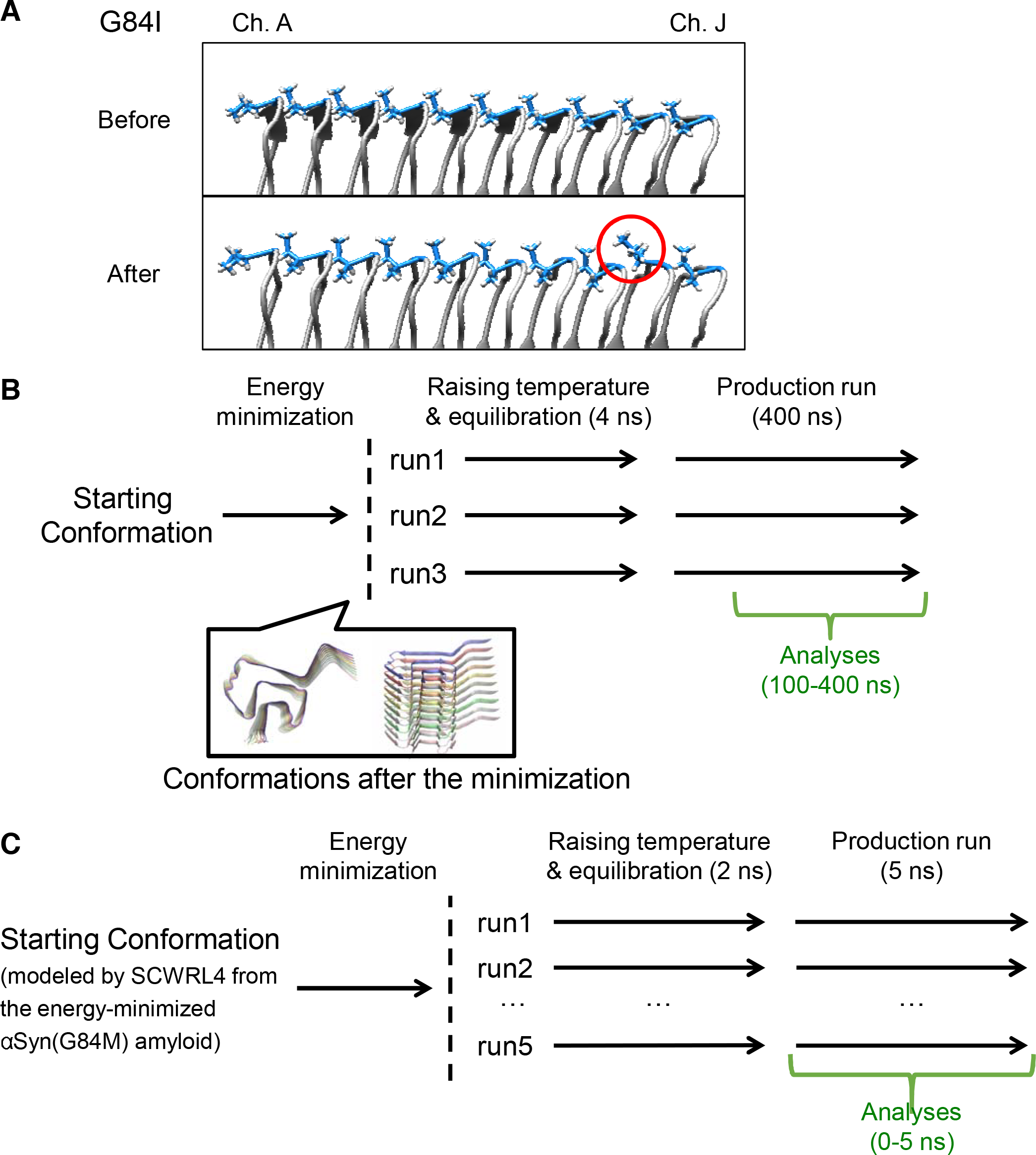
**A.** Side views of αSyn(G84I) amyloids before and after the energy minimization. Note that a side chain of Ile84 is already flipped immediately after the minimization step (red circle). **B.** A schematic illustration of the time course of MD simulation of 400 ns. **C.** A schematic illustration of the time course of short MD simulation of 5 ns, based on the “energy-minimized” αSyn(G84M) mutant amyloid.

**Supplementary Figure S2.**
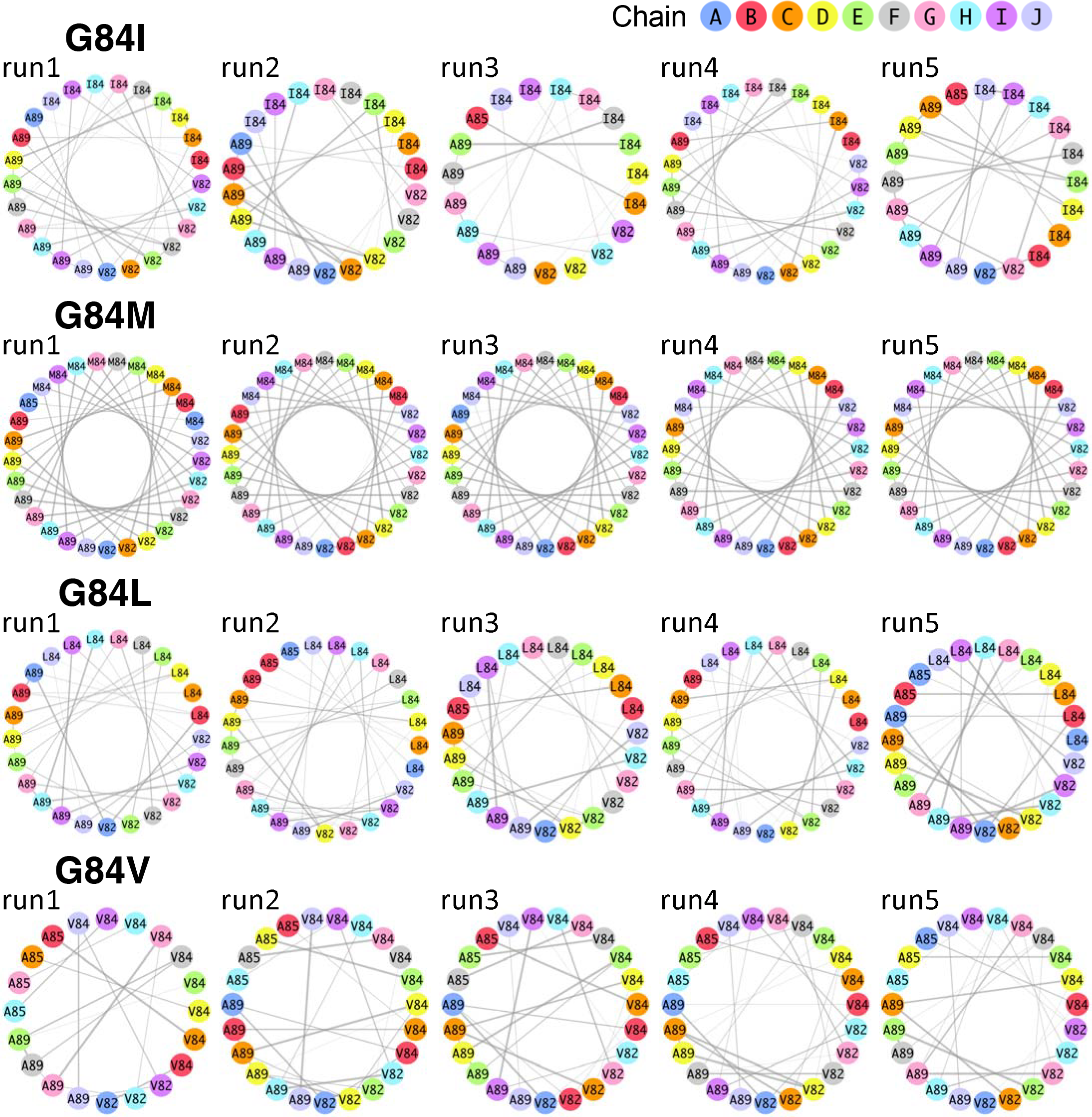
All the hydrophobic contact diagrams of αSyn(G84X) mutants calculated from the 5-ns short MD simulations.

